# The genetic legacy of continental scale admixture in Indian Austroasiatic speakers

**DOI:** 10.1101/423004

**Authors:** Kai Tätte, Luca Pagani, Ajai K. Pathak, Sulev Kõks, Binh Ho Duy, Xuan Dung Ho, Gazi Nurun Nahar Sultana, Mohd Istiaq Sharif, Md Asaduzzaman, Doron M. Behar, Yarin Hadid, Richard Villems, Gyaneshwer Chaubey, Toomas Kivisild, Mait Metspalu

## Abstract

Surrounded by speakers of Indo-European, Dravidian and Tibeto-Burman languages, around 11 million Munda (a branch of Austroasiatic language family) speakers live in the densely populated and genetically diverse South Asia. Their genetic makeup holds components characteristic of South Asians as well as Southeast Asians. The admixture time between these components has been previously estimated on the basis of archaeology, linguistics and uniparental markers. Using genome-wide genotype data of 102 Munda speakers and contextual data from South and Southeast Asia, we retrieved admixture dates between 2000 – 3800 years ago for different populations of Munda. The best modern proxies for the source populations for the admixture with proportions 0.78/0.22 are Lao people from Laos and Dravidian speakers from Kerala in India, while the South Asian population(s), with whom the incoming Southeast Asians intermixed, had a smaller proportion of West Eurasian component than contemporary proxies. Somewhat surprisingly Malaysian Peninsular tribes rather than the geographically closer Austroasiatic languages speakers like Vietnamese and Cambodians show highest sharing of IBD segments with the Munda. In addition, we affirmed that the grouping of the Munda speakers into North and South Munda based on linguistics is in concordance with genome-wide data.

## Introduction

Genetically diverse^1–3^ South Asia is home to more than a billion people who belong to thousands of distinct socio-culturally or ethnically defined population groups. These groups speak languages of four major language families: Indo-European, Dravidian, Austroasiatic and Trans-Himalayan. Studies based on genome-wide genotype data have shown that the majority of present day populations of the Indian subcontinent derive their genetic ancestry to a large extent from two ancestral populations – ancestral northern and southern Indians – of which the former is genetically close to West Eurasian populations^4–6^. In addition to these two components, the Munda speakers of the Austroasiatic family share a minor proportion of their genetic ancestry with Southeast Asian populations^7^. Austroasiatic languages are spoken by more than 100 million people in Mainland Southeast Asia (MSEA) and >10 million Austroasiatic speakers^8^ of Munda languages live in East and Central parts of India where they are surrounded by Indo-European, Dravidian and Trans-Himalayan languages speakers.

Considering the widespread sharing of words related to rice agriculture in all main branches of Austroasiatic, it has been proposed that this language family co-expanded with farming in MSEA and that the speakers of Munda languages spread to India as part of this farming expansion^9,10^. Alternatively, considering the deep splits of extant Munda and extinct Para-Munda languages and evidence for independent domestication of rice in India and in Southeast Asia, it has been proposed that Austroasiatic languages could have, instead, spread from India to Southeast Asia^11^. Given that about 25% of the genetic ancestry of Munda speakers has been shown to be shared with Southeast Asians, unlike in other Indian populations, and, reversely, because Austroasiatic speakers of Myanmar share some ancestry (∼16%) with Indian populations, it has been proposed that the expansion of rice farming may have involved bilateral movement of people^7^.

Studies analysing mtDNA and Y chromosome markers have revealed a sex-specific admixture pattern of admixture of Southeast and South Asian ancestry components for Munda speakers. While close to 100% of mtDNA lineages present in Mundas match those in other Indian populations, around 65% of their paternal genetic heritage is more closely related to Southeast Asian than South Asian variation^7,12,13^. Such a contrasting distribution of maternal and paternal lineages among the Munda speakers is a classic example of ‘father tongue hypothesis’^14^. However, the temporality of this expansion is contentious^7,13,15,16^. Based on Y-STR data the coalescent time of Indian O2a-M95 haplogroup was estimated to be >10 KYA^7,13^. Recently, the reconstructed phylogeny of 8.8 Mb region of Y chromosome data showed that Indian O2a-M95 lineages coalesce within a clade nested within East/Southeast Asian within the last ∼5-7 KYA^17^. This date estimate sets the upper boundary for the main episode of gene flow of Y chromosomes from Southeast Asia to India.

Previous autosomal study was limited to a single Austroasiatic population from Southeast Asia^7^, therefore in the present study, we generated and assembled large body of contextual genome-wide genotype data from Southeast Asia as well as from South Asia (Supplementary Table S1). We set out to affirm the signal of the admixture event in autosomal data and to address previously unresolved questions including: i) autosomal date of the South and Southeast Asian admixture event in Munda; ii) characteristics of the Indian ancestry component of the Mundas; iii) who are the closest living descendants of the source populations of the ancient admixture; iv) and if the grouping of the Munda speakers into North and South Mundas based on some linguistic models is supported by genetic data.

To address these questions, we analysed 102 individual samples from Munda speaking populations (including 10 newly reported samples) in context of 978 other samples (including 46 newly reported samples) from 72 populations mainly from India, Southeast Asia and East Asia. The Munda speakers are divided into North Mundas (NM) and South Mundas (SM) based on linguistic affinities. List of all the populations, sample sizes, and some additional information on the dataset can be found in Supplementary Table S1.

## Results and Discussion

### The Munda speakers as an admixed population

We first analysed Munda genomes with ADMIXTURE and PCA in context of other South and Southeast Asian populations and found that Munda share about three quarters of their genetic ancestry (k3 – k5 components in Figure 1) with Indian Dravidian and Indo-European speakers. Interestingly, Indian populations with the k3-k5 components have also a pink component (k2) which is widespread in European, West Eurasian, Near Eastern and Pakistani populations but missing in the Munda speakers. Roughly one quarter of the ancestral components in the Mundas’ genome (k6 – k12) are shared with Southeast Asians. There are two populations with a similar genetic profile to the Mundas in Central India: Dravidian speaking Gond who are known to have received a substantial gene flow from the Munda speakers^18^ and a linguistic isolate Nihali.

**Figure 1.**
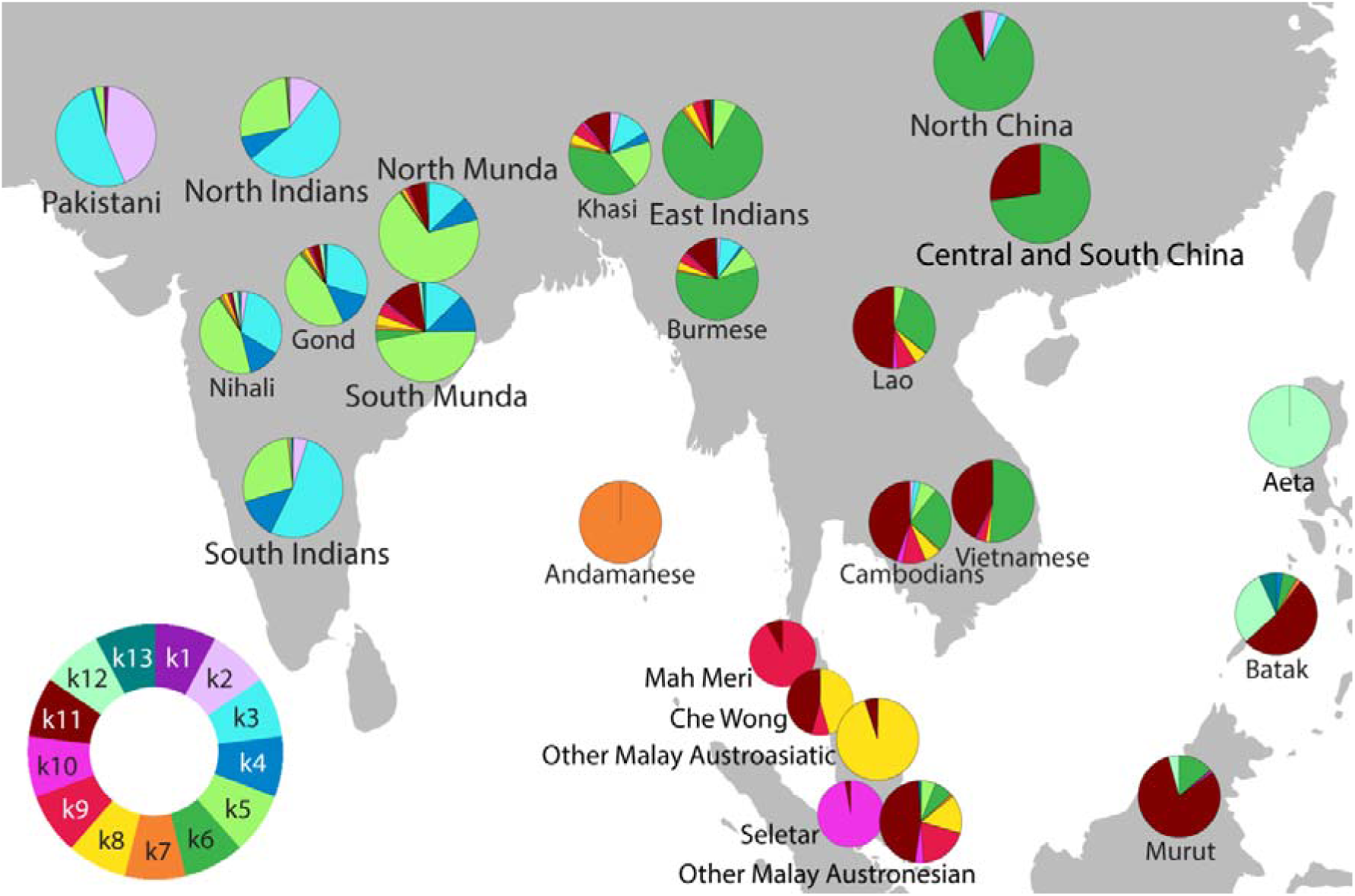
The distribution of genetic components (K=13) based on the global ADMIXTURE analysis (Supplementary Figure S1, S2, S3) for a subset of populations on a map of South and Southeast Asia. The circular legend in the bottom left corner shows the ancestral components corresponding to the colours on pie charts. The sector sizes correspond to population median.

Principal component analysis (PCA) roughly reflects geographical locations of studied populations (see Supplementary Fig. S4). Based on the first two components of PCA, the Mundas are genetically situated between South Asians and Southeast Asians and Oceanians. Furthermore, South and North Munda tribes are clearly different – South Mundas are genetically closer to Southeast Asians and Oceanians while North Mundas are closer to South Asians. In sum, the results of the ADMIXTURE and PCA are consistent with the model by which the genetic ancestry of Indian Munda speakers represents an admixture between Indian and Southeast Asian ancestries.

The scenario of independent evolution without admixture was rejected by 3-population formal test of admixture^6^ for South Munda, Santhal (NM) and Ho (NM) speakers, as they yielded significantly negative f3 values (indicative of admixture) when tested together with populations from India and Southeast Asia (Supplementary Table S2). Birhor (NM) and Korwa (NM) speakers did not display significant admixture signal potentially because of the vast genetic drift they have gone through after the admixture event as they show the lowest average heterozygosity among the Munda speakers (Supplementary Table S3). To understand further the position of Mundas in the genetic landscape of Indian populations, we plotted the second and third principal components from the global PCA analysis (see Supplementary Fig. S5). The Mundas were situated close to the Dravidian speaking southern Indian end of the gradient, near Pulliyar population from southwestern India, being stretched towards Southeast Asian populations, the closest ones being Bateq, Jehai, Kintaq and Mendriq from Malaysia.

### The best contemporary proxies for admixture sources

Three populations that yield the highest outgroup-f3 values as sources of Southeast Asian ancestry in Munda are Lao from Laos, Dai from China and Murut from Borneo. From South Asia, the populations that produce the highest f3 scores are Dravidian speaking Paniya and Pulliyar from Kerala region of India. For North Mundas, among the top Indian populations is also Indo-European speaking Chamar, whereas for South Mundas, there are Jarawa and Onge from Andaman Islands (Supplementary Table S2). Consistently, the South Munda speakers are the biggest DNA chunk donors from India to the Andamanese populations based of fineSTRUCTURE^19^ analysis (see Supplementary Fig. S7).

For a more detailed view of the temporary aspects of admixture, we assessed the sharing of DNA segments that are identical by decent between Munda speakers and other populations. Refined IBD analysis^20^ showed that from India, Mundas share the highest number of DNA segments identical by descent (IBD) with Dravidian speaking Chenchus (1.68; CI: 1.46 – 1.91) and Indo-European speaking Chamar (1.63; CI: 1.26 – 2.11) when disregarding Nihali and Gond tribes as Nihali, a language isolate, are possibly related to Munda and the Gond are reported to have received gene flow from the Mundas^18^. From Southeast Asia the sharing is highest with Mah Meri (2.04; CI: 1.79 – 2.33) and Temuan (1.93; CI: 1.67 – 2.24) tribes from Peninsular Malaysia, followed by Jakun and Che Wong from the same area (Figure 2, Supplementary Table S3). Surprisingly, the geographically closer Austroasiatic speakers from Southeast Asia, such as Cambodians and Vietnamese, do not share as many IBD segments with the Mundas. This effect could be caused by the fact that the mainland Southeast Asian populations have smaller proportions of the original Austroasiatic component in their genomes due to subsequent gene flow received from East Asia. Another explanation could be a more complex direction of gene flow in this area. Similar results were observed when using total lengths of shared IBD segments instead of their counts (Supplementary Figure S9).

**Figure 2.**
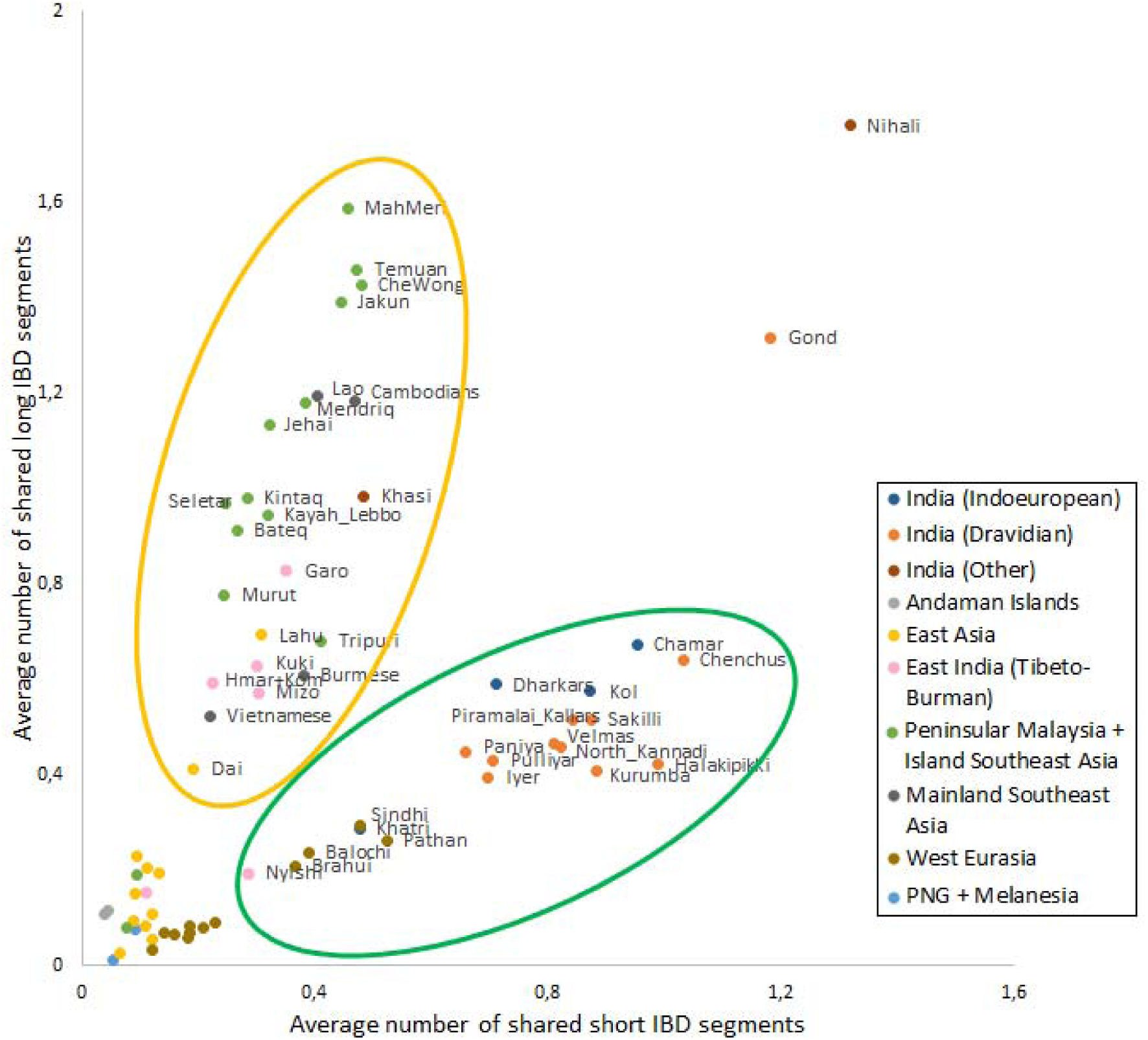
The plotted average counts of IBD segments up to 1 cM (short) and over 1 cM (long) shared with the Munda speakers. The points are coloured based on linguistics and geography according to the legend on the right.

When dividing the segments shared with the Mundas into two groups, short (<1 cM) and long (>1 cM), we noticed that the two sources, South Asian and Southeast Asian populations, clearly form two distinct groups based on shared segment length patterns (Figure 2). Both, mainland and island Southeast Asian populations share a high number of long IBD segments with the Mundas while Indian Dravidian and Indo-European speaking populations share plenty of short IBD segments. Surprisingly, no difference was found in Indian Dravidian and Indo-European speakers in context of sharing DNA with the Mundas (Welch’s t-test; short IBD *P* = 0.5218; long IBD *P* = 0.5302; all IBD *P* = 0.9305). The formation of the two groups seen on Figure 2 could refer to different genetic distance between admixed populations and other populations from the corresponding areas; *i.e.*, the Southeast Asian share of the Munda speakers’ genomes has diverged from present day Southeast Asians more recently than the South Asian part from present day South Asians. This result has to be taken with caution as we found correlation between the shared IBD segment lengths and the average heterozygosity in these populations (Supplementary Figure S8, Supplementary Table S3).

### Admixture proportions suggest a novel scenario

We used qpAdm^21^ to determine the relative proportions of West, Southeast and South Asian ancestries in Munda speakers, using a number of modern and ancient West Asian populations, Lao, and Onge or Paniya as proxies for the three Asian components (Supplementary Table S4). Regardless of which West Asian population we used, we found that Munda speakers can be described on average as a mixture of ∼19% Southeast Asian, 15% West Asian and 66% Onge (South Asian) components. Alternatively, the West and South Asian components of Munda could be modelled using a single South Asian population (Paniya), accounting on average to 77% of the Munda genome. When rescaling the West and South Asian (Onge) components to 1 to explore the Munda genetic composition prior to the introduction of the Southeast Asian component, we note that the West Asian component is lower (∼19%) in Munda compared to Paniya (27%) (Supplementary Table S4: *Average_Lao=0). Consistently with qpGraph analyses in Narasimhan et al. (2018)^22^, this may point to an initial admixture of a Southeast Asian substrate with a South Asian substrate free of any West Asian component, followed by the encounter of the resulting admixed population with a Paniya-like population. Such a scenario would imply an inverse relationship between the Southeast and West Asian relative proportions in Munda or, in other words, the increase of Southeast Asian component should cause a greater reduction of the West Asian compared to the reduction in the South Asian component in Munda. However, we note that the scaled proportion of West and South Asian components in our North and South Munda are comparable (Supplementary Table S4: Average_SM_Lao=0 and Average_NM_Lao=0 both show ∼18% West Asian and ∼82% South Asian contributions) while the Southeast Asian component is higher in South than in North Munda. The independence between the amount of Southeast and West Asian components in our North and South Munda populations contradicts the expectations and therefore points to an opposite and simpler scenario: both South and North Munda could be modelled as an initial admixture between Southeast Asian populations and an autochthonous Indian group with a slightly lower West/South Asian composition compared to what observed in Paniya today. South Munda then kept isolated from additional gene flow, while North Munda received a longer admixture pulse from the local Indian groups, which caused the dilution of the newly arrived Southeast Asian components in North Munda, without affecting the relative proportions of West and South Asian components.

### Dating the admixture event

We used ALDER to test this scenario and to infer the admixture time that led to the genesis of the Mundas^23^. The admixture midpoint was 3846 (3235 – 4457) years ago for South Mundas, which may point to the time of arrival of the Southeast Asian component in the area, and 2867 (1751 – 4525) years ago for North Mundas (Figure 3). The longer (1000 years) admixture time between North Munda and local Indian populations is consistent with the ADMIXTURE, PCA and qpAdm results where we saw North Mundas having a bigger proportion of Indian ancestry (made up, proportionally, by ∼18% West and 82% South Asian) and a smaller Southeast Asian fraction than South Mundas (Supplementary Figure S3, Supplementary Figure S4, Supplementary Table S4).

**Figure 3.**
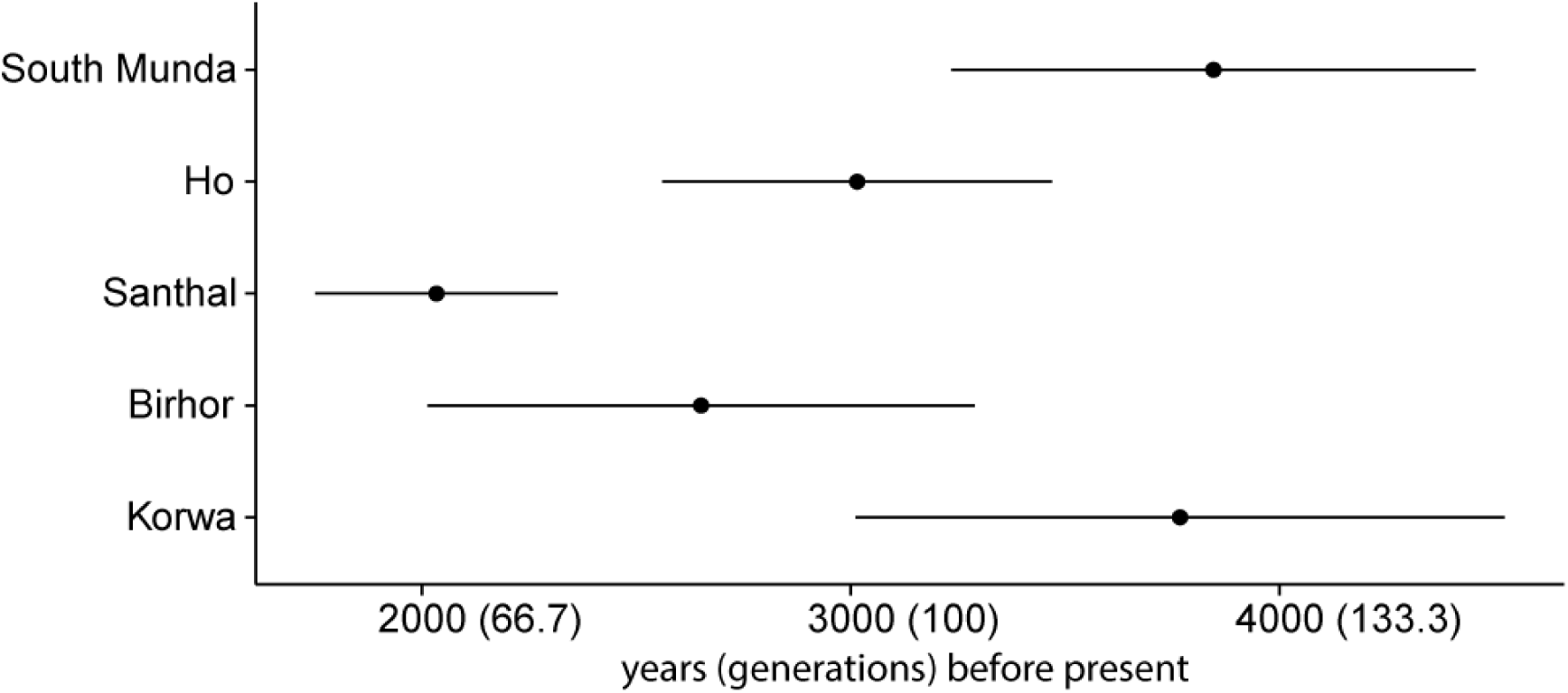
Admixture times as evaluated by ALDER. We let ALDER pair up populations from Southeast Asia and South Asia as several populations from either area were good proxies for the admixture event based on Refined IBD and f3 analyses. For accuracy, North Munda speaking Santhal, Ho, Korwa and Birhor were addressed separately as admixed populations; due to a small sample size South Munda speakers were treated as one population. Reference population pair was chosen based on LD decay curve amplitude. Standard errors are estimated by jackknifing on chromosomes. Generation length is 30 years^45^. For all the pairs, see Supplementary Table S5.

While the ALDER dates that we obtained are, to our knowledge, the first estimates of the time of admixture of the Munda speakers based on genome-wide data, estimates from previous studies, based on other types of data, have yielded much earlier dates for the spread of Austroasiatic populations in India. Diamond and Bellwood^24^ have estimated the age of the Munda speakers and cultivation of rice in India 5000 years old based on archaeological data. The Munda branch split from other Austroasiatic languages less than 7000 years ago based on Fuller’s archeolinguistic reconstruction^11,25^. Recent Y chromosome studies, based on large scale resequencing of the whole Y chromosome, have estimated the age of haplogroup O2a, in which the East Asia component of the Munda Y chromosomes is nested within, to much more recent dates than the earlier estimates based on short tandem repeat variation^7^. The entire Southeast Asian Y chromosome variation within the clade O2a2 has been estimated to be only 5 965 (CI 5 312 – 7 013) years old^17^, while the variation within Munda speakers has been estimated to derive from a single male ancestor who lived 4 300 (+- 200) years ago^15^. The latter date estimate is very similar to ours and implies a significant male-specific founder event as part of the admixture process.

In this study, we have replicated a result previously reported in Chaubey et al. (2011)^7^ that the Mundas lack one ancestral component (k2) that is characteristic to Indian Indo-European and Dravidian speaking populations. If this component came to India through one of the Indo-Aryan migrations^28^ then it would be fair to presume that the Munda admixture happened before this component reached India or at least before it spread all over the country. However, the admixture time computed here, falls in the exact same timeframe as the ANI-ASI mixture has been estimated to have happened in India^5^ through which the k2 component probably spread. Therefore, we propose that if the Munda admixture happened at the same time, it is possible for it to have happened in the eastern part of the country, east of Bangladesh, and later when populations from East Asia moved to the area, the Mundas migrated towards central India. Such a scenario, which may be further clarified by ancient DNA analyses, seems to be further supported by the fact that Mundas harbor a smaller fraction of West Asian ancestry compared to contemporary Paniya (Supplementary Table S4) and cannot therefore be seen as a simple admixture product of Southern Indian populations with incoming Southeast Asian ancestries.

### Sex-biased admixture in Munda speakers

In Chaubey et al. (2011)^7^, it was shown that the Munda speakers have high frequencies (19-95%) of East Asian chromosome Y haplogroup O2a at the background of almost no detectable East Asian mitochondrial DNA signal pointing to a sex-biased nature of admixture between Austroasiatic speakers and their local Indian neighbouring populations. We used outgroup f3 analysis to contrast allele frequency patterns on the X chromosome versus those on the autosomal chromosomes to clarify the maternal side of this sex-biased admixture event. Our analysis revealed that on X chromosome, a Dravidian speaking group, North Kannadi, is relatively more similar to Munda speakers than on autosomes, while on autosomes Lao, Vietnamese and Burmese from Southeast Asia and Sino-Tibetan speaking Kuki from India have relatively higher f3 values than on X chromosome (Supplementary Figure S12). This relatively higher autosomal affinity to Southeast Asian populations, however, is detectable only when testing South Munda speakers. The fact that South Munda speakers show more evident signs of a sex-specific admixture on maternal side is in accordance with the Y chromosome results from Chaubey et al. (2011), where South Munda speakers have also higher (0.73) average frequency of haplogroup O2a than North Munda speakers (0.62)^7^. This finding is consistent with our proposed scenario where South Munda kept isolated after the admixture event, while North Munda received additional admixture from local Indian groups, which diluted Southeast Asian component and blurred the signs of the sex-specific nature of the admixture event as the latter admixture pulse in North Munda was not sex-specific anymore.

### Linguistics is in concordance with genome-wide data

Until now, we have presumed that the linguistic classification of the Mundas (North and South) is a suitable grouping criteria for genetic analyses. Here we take a glance at the genetic relationship between different North and South Munda populations. PCA of only Munda populations displayed North and South Mundas as separate groups, except one Juang and one Kharia individual fell together with North Mundas on first two principal components (see Supplementary Fig. S6). ADMIXTURE analysis showed that North Mundas have less of the combined k8 – k11 genetic component than South Mundas (Wilcoxon rank sum test; N1 = 75; N2 = 11; *P* < 0.0001). These components were maximised in East and Southeast Asian samples. Smaller amount of Lao ancestry in North Mundas was also shown by qpAdm analysis (Supplementary Table S4). On the fineSTRUCTURE tree^19^, North and South Mundas clustered separately, except Kharia samples (South Munda) which clustered with Asur and Ho samples from North Munda (Figure 4). All these analyses showed that Kharia and Juang were the most similar population to North Mundas among South Munda populations. Refined IBD analysis infers that North Munda populations share more long and short IBD segments among each other than with South Munda populations (see Supplementary Fig. S10). Therefore, by and large, the linguistic classification justifies itself but Kharia and Juang do not fit in this simplification perfectly. Interestingly, although Diffloth’s classification of the Munda languages into North and South Munda^26^ is widely cited, in 2005, Diffloth changed the position of Kharia-Juang branch on the language tree from South Munda group to be a side branch of the group that was previously known as North Munda^27^. Hence, this is in accordance with our findings about Juang and Kharia genetic affinities.

**Figure 4.**
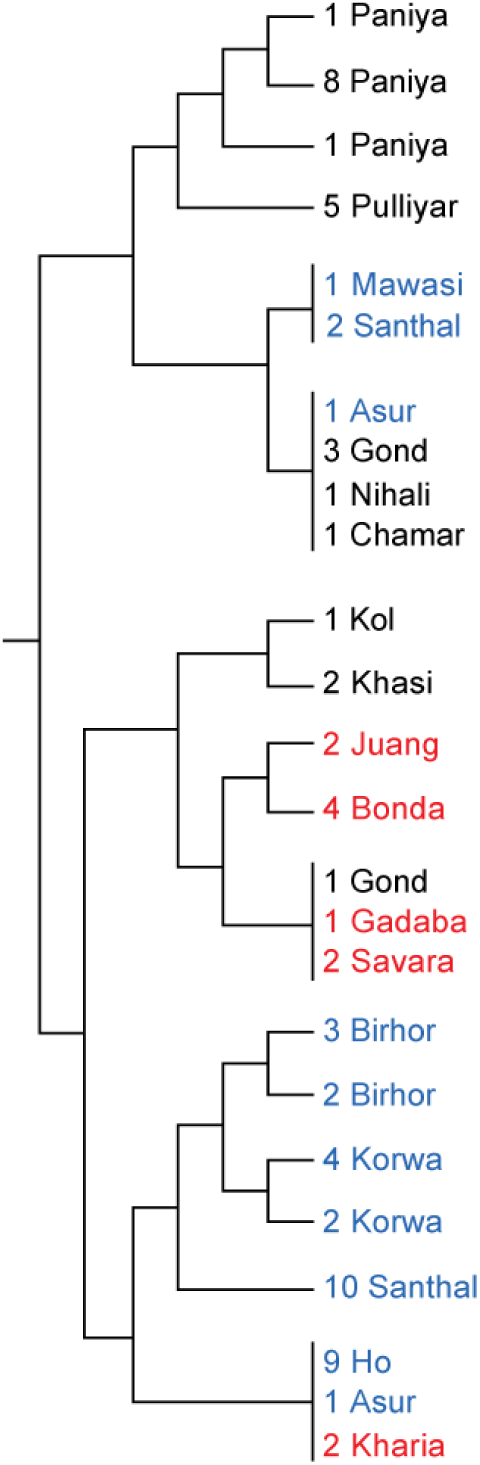
A branch from a FineSTRUCTURE tree where all the Munda samples used in this analysis are situated on. Samples are coloured as follows: North Munda speakers – blue, South Munda speakers – red.

## Methods

### Samples Collection and Genotyping

The analyses were performed on a merged dataset of 56 new samples together with 1024 previously published samples from different studies^4,7,29–37^ (Supplementary Table S1). The new samples were collected from Laos (Lao N = 24), Bangladesh (Santhal (NM) N = 10), and East India (Hmar N = 4, Kom N = 2, Kuki N = 6, Mizo N = 5, Naga N = 1, Nyishi N = 4). DNA was extracted from blood samples collected from healthy adult donors who signed an informed consent form. New samples were genotyped using Illumina OmniExpress Bead Chips for 730k, 710k and 650k SNPs. The study was approved by Research Ethics Committee of the University of Tartu. All genotyped data will be made publicly available on the ebc.ee/free_data website.

### Data Curation

All the samples were filtered with plink v1.9^38^. Only SNPs on autosomal chromosomes with a minor allele frequency > 1% and genotyping success > 97% were used in the analyses. Only individuals with a genotyping success rate > 97% were left in the sample set. 245848 variants and 1072 people passed the filters; 8 Gond were removed due to low genotyping success rate. For analyses that are affected by linkage disequilibrium (PCA, ADMIXTURE), dataset was further pruned by excluding SNPs with pairwise genotypic correlation r^2^ > 0.4 in a window of 200 SNPs sliding the window by 25 SNPs at a time^39^. This left us 155743 SNPs.

### Population Structure Analyses

To capture genetic variability, we performed PCA using software EIGENSOFT 6.1.4^40^ on pruned data of the whole filtered dataset (1072 individuals). To get some idea of the Munda speakers’ genetic structure in context of other Asian populations, we ran ADMIXTURE 1.23 program^41^ with random seed number generator on the LD pruned data set one hundred times at K = 2 to K = 18 (Supplementary Figure S1). Following an established procedure, we examined the log likelihood scores (LLs) of the individual runs and found that the highest K with stable (global maximum has been reached) LL values is K = 13. Based on cross-validation (CV) procedure, genetic structure of a sample set is best described choosing the value of K with the lowest CV error. In our dataset the lowest CV error was at K = 13 (Supplementary Figure S2).

### Tests Aimed at Providing Demographic Inferences

To test the admixture, we ran three-population formal test of admixture^6^ using popstats program by Skoglund et al. (2015)^42^. For f3 analysis, source 1 was South Asian or West Eurasian population and source 2 was Southeast Asian or East Asian population. Outcomes with |Z| > 3 were considered significant. All the South Munda speaking tribes (Bonda, Gadaba, Juang, Kharia, Savara) were treated as one population due to small sample size. We ran outgroup f3 statistic as f3 = (SouthMunda/Ho(NM), X, Yoruba) to find the closest modern populations from out data set for South and North Munda.

To retrieve the admixture proportions, we run the qpAdm software^21^ testing the following South and North Munda populations (Bonda, Gadaba, Juang, Kharia, Savara, Asur, Birhor, Ho, Korwa, Mawasi, Santhal) as a three ways mixture of all possible combinations of West (Anatolia_N, Armenia_MLBA, Germans, Iran_N, IranianLaz2016), East (Lao) and South (Onge, Paniya) Asian groups and using as outgroups the following groups (Natufian, WHG, Han, Kankanaey, Karitiana, MbutiLaz2016, Papuan, Ust_Ishim, Yorubas)^43,44^.

We used ALDER^23^ to infer admixture dates for South Munda, Ho (NM), Santhal (NM), Birhor (NM) and Korwa (NM). We used all the populations spanning from India to Europe from our data set as source 1 and all the populations from East and Southeast Asia as source 2. The population pairs to represent admixture times were chosen based on decay status and LD decay curve amplitude. Standard errors were estimated by jackknifing on chromosomes. We used generation length of 30 years^45^.

### Haplotype-based Analyses

To investigate the relationship between the Munda speakers and Andmanese, we used fineSTRUCTURE^19^. For this analysis, the data was previously phased with Beagle 3.3.2^46^. A co-ancestry matrix was constructed using ChromoPainter v1^19^ with the default settings. From the co-ancestry matrix, the mean chunk lengths donated by Eurasian populations to Jarawa and Onge were extracted. Beagle was also used in Refined IBD^20^ analysis, where we studied the sharing of DNA segments of identity-by-descent (IBD) between the Munda speakers and other populations in our data set. From the results, we extracted the count of segments shared between every two individuals and found population medians. We did the same with short (<1 cM) and long (>1 cM) segments, to find patterns. We also compared total length of IBD segments shared between individuals from two different populations on average.

All the methods were performed in accordance with relevant guidelines and regulations.

## Acknowledgments

Support was provided by the European Union through the European Regional Development Fund projects i) Centre of Excellence for Genomics and Translational Medicine Project No. 2014–2020.4.01.15-0012 (K.T., M.M., T.K.) and ii) Project No. 2014-2020.4.01.16-0024, MOBTT53 (L.P.) and Estonian Institutional Research grants IUT 24-1 (K.T., T.K., A.P., D.M.B., M.M., and R.V.) and IUT 20-46 (S.K.). G.C. was supported by National Geographic explore grant HJ3-182R-18.

## Author Contributions

M.M., T.K. and G.C. devised and supervised the study. K.T. wrote the manuscript with input from M.M., T.K., L.P., G.C. and A.P. G.C., S.K., B.H.D., X.D.H., D.M.B., Y.H., G.N.N.S., M.I.S., M.A., R.V. and M.M. performed anthropological work, sample collection and provided laboratory and computing facilities. Data analyses were performed by K.T., G.C. and L.P. Figures were prepared by K.T. All authors have reviewed the manuscript.

## Additional Information

The authors declare no competing interests.

## References

1. Mallick, S. et al. The Simons Genome Diversity Project: 300 genomes from 142 diverse populations. Nature 538, 201–206 (2016).

2. Pagani, L. et al. Genomic analyses inform on migration events during the peopling of Eurasia. Nature 538, 238–242 (2016).

3. Xing, J. et al. Genetic diversity in India and the inference of Eurasian population expansion. Genome Biol. 11, R113 (2010).

4. Metspalu, M. et al. Shared and Unique Components of Human Population Structure and Genome-Wide Signals of Positive Selection in South Asia. Am. J. Hum. Genet. 89, 731–744 (2011).

5. Moorjani, P. et al. Genetic Evidence for Recent Population Mixture in India. Am. J. Hum. Genet. 93, 422–438 (2013).

6. Reich, D., Thangaraj, K., Patterson, N., Price, A. L. & Singh, L. Reconstructing Indian population history. Nature 461, 489–494 (2009).

7. Chaubey, G. et al. Population Genetic Structure in Indian Austroasiatic Speakers: The Role of Landscape Barriers and Sex-Specific Admixture. Mol. Biol. Evol. 28, 1013–1024 (2011).

8. Census of India: Abstract of speakers’ strength of languages and mother tongues –2001. Available at: http://www.censusindia.gov.in/Census_Data_2001/Census_Data_Online/Language/Statement1.aspx. (Accessed: 3rd May 2018)

9. Bellwood, P. First farmers. Orig. Agric. Soc. (2005).

10. Higham, C. Chapter 18 Languages and Farming Dispersals: Austroasiatic Languages and Rice Cultivation. (2003).

11. Fuller, D. Q. Non-human genetics, agricultural origins and historical linguistics in South Asia. in The Evolution and History of Human Populations in South Asia 393–443 (Springer, Dordrecht, 2007). doi:10.1007/1-4020-5562-5_18

12. Kumar, V. et al. Y-chromosome evidence suggests a common paternal heritage of Austro-Asiatic populations. BMC Evol. Biol. 7, 47 (2007).

13. Zhang, X. et al. Y-chromosome diversity suggests southern origin and Paleolithic backwave migration of Austro-Asiatic speakers from eastern Asia to the Indian subcontinent. Sci. Rep. 5, (2015).

14. Forster, P. & Renfrew, C. Mother Tongue and Y Chromosomes. Science 333, 1390–1391 (2011).

15. Arunkumar, G. et al. A late Neolithic expansion of Y chromosomal haplogroup O2a1-M95 from east to west: Late Neolithic expansion of O2a1-M95. J. Syst. Evol. 53, 546–560 (2015).

16. Riccio, M. E. et al. The Austroasiatic Munda population from India and its enigmatic origin: a HLA diversity study. Hum. Biol. 83, 405–435 (2011).

17. Karmin, M. et al. A recent bottleneck of Y chromosome diversity coincides with a global change in culture. Genome Res. 25, 459–466 (2015).

18. Chaubey, G. et al. Reconstructing the population history of the largest tribe of India: the Dravidian speaking Gond. Eur. J. Hum. Genet. 25, 493–498 (2017).

19. Lawson, D. J., Hellenthal, G., Myers, S. & Falush, D. Inference of Population Structure using Dense Haplotype Data. PLOS Genet. 8, e1002453 (2012).

20. Browning, B. L. & Browning, S. R. Improving the Accuracy and Efficiency of Identity-by-Descent Detection in Population Data. Genetics 194, 459–471 (2013).

21. Patterson, N. et al. Ancient Admixture in Human History. Genetics 192, 1065–1093 (2012).

22. Narasimhan, V. M. et al. The Genomic Formation of South and Central Asia. bioRxiv (2018). doi:10.1101/292581

23. Loh, P.-R. et al. Inferring Admixture Histories of Human Populations Using Linkage Disequilibrium. Genetics 193, 1233–1254 (2013).

24. Diamond, J. Farmers and Their Languages: The First Expansions. Science 300, 597–603 (2003).

25. Fuller, D. Q. An agricultural perspective on Dravidian Historical Linguistics: Archaeological crop packages, livestock and Dravidian crop vocabulary. in Assessing the Languaging/Farming Dispersal Hypothesis (eds. Bellwood, P. & Renfrew, C.) 191–213 (McDonald Institute for Archaeological Research, 2003).

26. Diffloth, G. & Zide, N. Austro-asiatic languages. Encycl. Br. 2, 480–484 (1974).

27. Sagart, L., Blench, R. & Sanchez-Mazas, A. The peopling of East Asia putting together archaeology, linguistics and genetics. (RoutledgeCurzon, 2005).

28. Silva, M. et al. A genetic chronology for the Indian Subcontinent points to heavily sex-biased dispersals. BMC Evol. Biol. 17, (2017).

29. Aghakhanian, F. et al. Unravelling the Genetic History of Negritos and Indigenous Populations of Southeast Asia. Genome Biol. Evol. 7, 1206–1215 (2015).

30. Basu, A., Sarkar-Roy, N. & Majumder, P. P. Genomic reconstruction of the history of extant populations of India reveals five distinct ancestral components and a complex structure. Proc. Natl. Acad. Sci. 113, 1594–1599 (2016).

31. Behar, D. M. et al. The genome-wide structure of the Jewish people. Nature 466, 238–242 (2010).

32. Li, J. Z. et al. Worldwide human relationships inferred from genome-wide patterns of variation. Science 319, 1100–1104 (2008).

33. Migliano, A. B. et al. Evolution of the pygmy phenotype: evidence of positive selection fro genome-wide scans in African, Asian, and Melanesian pygmies. Hum. Biol. 85, 251–284 (2013).

34. Mörseburg, A. et al. Multi-layered population structure in Island Southeast Asians. Eur. J. Hum. Genet. 24, 1605–1611 (2016).

35. Pierron, D. et al. Genome-wide evidence of Austronesian-Bantu admixture and cultural reversion in a hunter-gatherer group of Madagascar. Proc. Natl. Acad. Sci. U. S. A. 111, 936–941 (2014).

36. Yunusbayev, B. et al. The Caucasus as an Asymmetric Semipermeable Barrier to Ancient Human Migrations. Mol. Biol. Evol. 29, 359–365 (2012).

37. Yunusbayev, B. et al. The Genetic Legacy of the Expansion of Turkic-Speaking Nomads across Eurasia. PLOS Genet. 11, e1005068 (2015).

38. PLINK 1.9. Available at: http://www.cog-genomics.org/plink/1.9/. (Accessed: 3rd May 2018)

39. Chang, C. C. et al. Second-generation PLINK: rising to the challenge of larger and richer datasets. GigaScience 4, 7 (2015).

40. Patterson, N., Price, A. L. & Reich, D. Population Structure and Eigenanalysis. PLOS Genet. 2, e190 (2006).

41. Alexander, D. H., Novembre, J. & Lange, K. Fast model-based estimation of ancestry in unrelated individuals. Genome Res. 19, 1655–1664 (2009).

42. Skoglund, P. et al. Genetic evidence for two founding populations of the Americas. Nature 525, 104–108 (2015).

43. Lazaridis, I. et al. Genomic insights into the origin of farming in the ancient Near East. Nature 536, 419–424 (2016).

44. Haak, W. et al. Massive migration from the steppe was a source for Indo-European languages in Europe. Nature 522, 207–211 (2015).

45. Fenner, J. N. Cross-cultural estimation of the human generation interval for use in genetics-based population divergence studies. Am. J. Phys. Anthropol. 128, 415–423 (2005).

46. Browning, S. R. & Browning, B. L. Rapid and Accurate Haplotype Phasing and Missing-Data Inference for Whole-Genome Association Studies By Use of Localized Haplotype Clustering. Am. J. Hum. Genet. 81, 1084–1097 (2007).

